# Conformational plasticity and sequence specificity interplay in non-canonical tandem RRM-RNA binding

**DOI:** 10.64898/2025.12.13.694091

**Authors:** Roxana Geanina Vasarhelyi, Vlad Cojocaru

**Affiliations:** Doctoral School for Integrative Biology, Faculty of Biology and Geology, Babeş-Bolyai University, Cluj-Napoca, Romania; Star-UBB Institute, Babeş-Bolyai University, Cluj-Napoca, Romania

**Keywords:** RRM domain, RNA-binding proteins, conformational change, sequence specificity, molecular recognition, tandem RRM, molecular dynamics simulation

## Abstract

The Dead End protein (DND1), a key regulator of germline cell fate, utilizes two RNA Recognition Motifs (RRM) in tandem to bind AU-rich RNA in a non-canonical manner. Only one RRM uses the known RNA binding interface, whereas the second motif has only minimal contacts with the RNA. To characterize the structural features and dynamics that contribute to RNA binding, we performed a series of atomistic molecular dynamics simulations and found that the complex is highly dynamic, deviating significantly from the experimental structure. We found that RNA binding restricts without abolishing the inter-domain motions of the RRMs and that cooperative binding of both RRMs is required to reduce the flexibility of the bound RNA. Through detailed analysis of the protein–RNA interactions, we show that the RNA binding interface in the tandem RRM–RNA complex remains stable despite extensive conformational plasticity. The RRMs cooperate to sustain a layered RNA binding mechanism, with aromatic stacking of the central adenosine with RRM1 residues as foundation, arginine residues anchoring the RNA backbone, and residues in the inter RRM linker forming hydrogen bonds with the RNA. Despite insufficient conformational space sampling to claim convergence, the protein-RNA interactions were consistent across multiple independent simulations with cognate and non-cognate RNA sequences. This substantiates our findings which showcase how structural dynamics impact RNA recognition, enabling structural adaptation for functional versatility in multi-domain proteins.

**Statement of Significance:** The RNA recognition motif (RRM) is the most abundant RNA binding domain in proteins. Often, one protein has multiple RRMs to enhance RNA binding specificity. For example, Dead End, an essential protein in germ cells with context dependent roles in cancer has two RRMs connected by a short linker. It is not understood how the inter-domain dynamics between the two RRMs modulate the RNA binding specificity. We found that the tandem RRMs of Dead End display extensive conformational plasticity in molecular dynamics simulations with variable inter-domain orientations. Despite this plasticity, the core RNA binding specificity is kept as a result of the cooperative binding of the two RRMs. Our findings provide evidence for highly dynamic and adaptable RNA recognition by tandem RRMs.

## Introduction

RNA-binding proteins (RBPs) are key regulators of gene expression. They interact with RNA to form ribonucleoprotein (RNP) complexes (1, 2). Most RBPs contain at least one RNA Binding Domain (RBD) (3), which allows them to control nearly all steps of RNA processing (4, 5). The RNA Recognition Motif (RRM) is the most common RBD in eukaryotes (2). Found in over 50% of human RBPs, with the human genome encoding 239 proteins that have at least one RRM domain (6). This prevalence highlights its critical role in post-transcriptional gene regulation by recognizing single-stranded RNA (ssRNA) (3).

The RRM fold is highly conserved (7), but it is also functionally versatile. Its core structure, established through high-resolution studies (8–12), is defined by a canonical β_1_α_1_β_2_β_3_α_2_β_4_ topology, consisting of four β-strands and two α-helices (3). Canonical RRM-RNA binding involves the central β sheet (3) and is controlled by two key consensus sequences: RNP1 on the β_3_ strand and RNP2 on the β1 strand (1, 3–5). These sequences bear aromatic amino acids which form sequence-specific interactions with the RNA bases (4). While a single RRM can bind up to eight nucleotides, multiple RRM domains often work together to increase binding specificity and affinity (13). Tandem RRM proteins exhibit diverse inter-domain orientations that determine their cooperative RNA binding mode, allowing the domains to either bind adjacent RNA sites or recognize sites separated by an RNA loop (14).

Some RRMs exhibit non-canonical features such as additional α-hairpins (15) or β-sheets (16, 17) For example, the Dead End protein (DND1) consists of two RRMs and a double-stranded RNA Binding Domain (dsRBD) (18, 19). As an essential, conserved RBP, DND1 is critical for maintaining primordial germ cell (PGC) fate and survival (19–21), where its loss is directly linked to the formation of germ cell tumors (22). Structural studies of its non-canonical RRMs (PDB-ID: 7Q4L) revealed an unusual binding mechanism for high-affinity recognition of an AU-rich consensus sequence (18). The two RRMs, RRM1 and RRM2, cooperate to achieve high-affinity binding to AU-rich RNA targets. RRM1 is the essential primary binding platform due to its extended RRM (eRRM) fold and, together with the ultra-short inter-domain linker, mediates the sequence-specific recognition of a central adenosine (A4) residue in the RNA. In contrast, the non-canonical RRM2, which lacks aromatic RNP residues and does not bind RNA alone, acts as a stabilizing lock. It achieves this by using an atypical binding pocket on its α-helix 2 to specifically recognize a uridine (U6) residue and cap the RNA, forcing an unusual 120° turn in the RNA backbone and increasing the overall binding affinity (18).

Functionally, DND1 serves as a dual regulator, capable of stabilizing target transcripts by blocking microRNA (miRNA) access (23), or conversely, promoting messenger RNA decay by recruiting a deadenylation complex, a mechanism controlled in part by the protein’s dsRBD (18, 19). These opposing regulatory functions are central to its involvement in disease, as discussed in a systematic review by Faraji et al. (22). DND1’s stabilizing role promotes anti-cancerous effects in some somatic cells, e.g. by stabilizing pro-apoptotic mRNA, whereas its repressive function is linked to tumor formation (18, 23). In development, DND1 partners with the NANOS3 (NANOS homolog 3) protein to suppress translation of genes such as SOX4, thereby constraining germ cell lineage specification (24). Conversely, a recent study showed that DND1 and NANOS3 also form a complex that activates the translation of specific RNAs in primordial germ cells by coupling 3′UTR binding to the translation initiation machinery (25). It revealed that the DND1-NANOS3 interaction can both repress and promote translation depending on the context.

Furthermore, while DND1 can act as a tumor suppressor in some somatic contexts (23), recent discoveries show a pro-tumorigenic role in melanoma, where the circular RNA binds directly to DND1, sequestering it and reversing its inhibitory effect on the PI3K/AKT (Phosphoinositide 3-kinase/protein kinase B) signaling pathway, thereby positioning the circular RNA as a potential diagnostic biomarker and therapeutic target (26). This functional versatility may require structural flexibility and adaptation.

Despite the availability of one experimental structure for the DND1-RNA complex (PDB-ID: 7Q4L), the role of conformational plasticity in this cooperative tandem RRM-RNA recognition mechanism remains unknown. We present here extensive molecular dynamics (MD) simulations of the DND1-RNA complex, the truncated complexes with each RRM bound to RNA and the apo protein and RNA. From these, we found that the complex is highly flexible, deviating significantly from the experimental structure. We show that RNA binding restricts but does not abolish the inter-domain motion of the two RRMs. At the same time, the cooperative tandem RRM-RNA binding restricts the conformational dynamics of the RNA. Despite the extensive conformational plasticity, the core RNA binding interface and sequence specificity is maintained in the tandem RRM-RNA complex. Analyzing the evolution of the protein-RNA interface during simulations, we found that the cooperative binding of the two RRMs is required for maintaining the interactions that define the core sequence specificity. We confirmed these findings with MD simulations with a mutated RNA in which the central adenine was replaced with an uracil. In agreement with previous experiments, the binding of the truncated RRM1 to RNA was more stable than the RRM2-RNA interaction. Our findings contribute to an intricate dynamics-centered mechanism of RNA recognition by multi-domain proteins which may allow structural adaptation for functional versatility.

## Methods

### Structure prediction with AlphaFold 3

We generated a total of 25 predicted structures for the protein-RNA complex by running 5 predictions using the AlphaFold 3 (AF3) server (27), each with a different seed number (resulting in five models per prediction). We selected the top five models from the complete set based on the highest pLDDT (predicted Local Distance Difference Test) scores, a measure of structural confidence. We assessed the quality and accuracy of these chosen models using a CAPRI-based (28) analysis performed in HADDOCK3 (29), using the complex’s NMR (nuclear magnetic resonance) ensemble (PDB-ID: 7Q4L) (18) as the reference experimental structure for validation. We quantified the accuracy of the predicted structures using the Interface Root Mean Square Deviation (IRMSD), which measures backbone atom deviation exclusively at the protein-RNA interface, and the Fraction of Native Contacts (FNAT), which reports the percentage of residue-residue contacts from the reference reproduced by the predicted structure. We used these metrics to quantify the deviation of the AF3 predicted structures from the experimental reference, noting that the PDB entry for the experimental structure was not within the AF3 training data.

### Modelling the systems for molecular dynamics simulations

From the experimental structure PDB-ID: 7Q4L (18) which contains an ensemble of 20 structures derived from NMR data, 3 frames were selected, each exhibiting slightly different conformations: two having the N-terminal extension (from G1 to N7) of RRM1 away from the globular domain but in different orientations, and one where it is closer to the RRM1 and the RNA binding site (see Fig. S1 for details). To better reflect the natural environment of the protein in which the RRMs are not isolated, we generated models of DND1 with extended terminal tails using Modeller (30). For this we added 10 amino acids to the N-terminal tail and 8 amino acids to the C-terminal tail (Fig S1A). We ran Modeller on the DND1 protein sequence (UniProt ID: Q8IYX4, Fig. S1A) (31) using the full 7Q4L NMR ensemble as template and selected the three best-scoring models. We superimposed them back to the original ensemble to identify the structural snapshots in the ensemble closest to each of them (Fig. S1B). To build the models, we used the selected structures from the 7Q4L ensemble as templates and aligned the DND1 protein sequence on it (UniProt ID: Q8IYX4).

To study the role of each RRM in RNA binding, we also built models of truncated systems which contain only one RRM bound to the RNA (RRM1-RNA and RRM2-RNA) simply by removing the other RRM. Furthermore, we prepared models of the unbound DND1 (tandem RRM) and unbound individual RRMs (RRM1 and RRM2) as well as the unbound RNA. Therefore, in total, we prepared 21 models for the follow-up MD simulations.

### Molecular dynamics simulations

We established seven molecular systems for simulations: the full DND1-RNA (RRM1-RRM2-RNA) complex, the free DND1 (RRM1-RRM2) tandem domain, the truncated RRM1-RNA and RRM2-RNA complexes, the free RRM1 and RRM2, and the free RNA. For each system, we performed three independent replicate simulations (M1, M2, and M3, named after the name of the starting model), including separate equilibration protocols (Table S1, see below). This resulted in a total of 21 simulations (Table S2). For the DND1-RNA complex started from the M2 model, one replica showed large structural changes during the equilibration step. Therefore, we performed two additional replicas starting from the M2 model for the DND1-RNA, RRM1-RNA, and RRM2-RNA systems. We also performed a separate set of simulations in which the central A4 nucleotide of the RNA was mutated to a uridine (A4U), as previous experimental work showed that the adenine at this position is essential for protein–RNA interaction. These simulations (3 replicas for each system started with the M1, M2, M3 models) were run for the DND1-RNA^A4U^ and RRM1-RNA^A4U^ systems.

We solvated all systems using the OPC water model (32) neutralized with counterions, Na⁺ or Cl⁻. We used 5 Å as the minimal buffer distance between the solute and the edge of the water box. This is shorter than the distance usually used but since we have extended the N-terminal and C-terminal regions, the box size is large enough for the symmetric molecules to not interact with each other. Subsequently, 100 mM NaCl and 100 mM KCl were added to reflect buffer conditions. We used the AMBER ff19SB force field (33) and the OL3 RNA force field (34), for the protein and RNA respectively. We minimized the energy of the solvated systems with the AMBER software (35). Equilibration was carried out under constant temperature and pressure using NAMD (36). The equilibration consisted of 17 sequential steps, totaling 4.875 ns (Table S1). The protocol systematically reduced positional restraints, beginning with restraints on all solute atoms and gradually removing them until the final unrestrained steps. Temperature was maintained at 300 K using Langevin dynamics with a damping coefficient of 1 ps⁻¹, and pressure was controlled at 1 atm using the Langevin piston method. Short-range nonbonded interactions were truncated at 9 Å cutoff, and long-range electrostatics were computed using the Particle Mesh Ewald (PME) method. MD simulations were performed in the isothermal-isobaric (NPT) ensemble for 1 μs with a timestep of 2 fs. The temperature was kept constant at 300 K using Langevin dynamics with a damping coefficient of 0.1 ps. The pressure was maintained at 1 atm using the Langevin Piston method, with a piston period of 1.5 ps and a decay time of 1.2 ps, respectively. All simulations were performed with NAMD. Snapshots were selected for analysis every 5 ps.

### Analysis of MD simulations

We processed and analyzed the MD simulations using CPPTRAJ (37). We first wrapped the molecules back into the central unit box, centering on the structural part. The added extended tails (residues 1-8 and 234-242) were wrapped but not used in the selection for defining the center of the box. Then, we generated trajectories and topology files by removing the water and the ions.

To quantify the conformational flexibility, we calculated the Root Mean Square Fluctuation (RMSF) using all heavy atoms (non-hydrogen atoms) and averaging per residue. The RMSF was determined for the trajectory of each replica (M1, M2, M3) individually, rather than by using a single ensemble of the three simulations/system. We calculated Root Mean Square Deviations (RMSD) using the minimized structure that was closest to the average coordinates of the 20 experimental NMR structures as reference. For all protein-containing systems, we performed the fit on the structural core of the protein (excluding the added extended tails) to remove translational and rotational motion. Depending on the question asked, the fit was performed either on RRM1 or RRM2, or both domains. For the simulation involving RNA, we performed the fit on the entire RNA molecule.

To capture the motion of the RRMs relative to each other and their compactness, we calculated the distance between the center of mass (COM-COM) of the two RRMs and their radius of gyration (ROG) using again only the structural regions of the domains.

To evaluate the accessibility of the protein-RNA interface, we calculated the surface accessible solvent area (SASA) using the surf command in CPPTRAJ for the β-sheets of RRM1 (residues 33-37, 38-44, 58-64, 85-90, 99-105) and RRM2 (residues 138-143, 165-172, 180-184, 210-214).

### Hierarchical Clustering

To analyze the different conformational states sampled during the simulations, we performed hieragglo (Hierarchical Agglomerative) clustering in CPPTRAJ (38) based on the RMSD of DND1 (RRM1-RRM2) upon fitting only RRM1. We reduced the size of the dataset by considering only one out of every three frames (33%) of the trajectories, resulting in an effective time interval of 15 ps between consecutive analyzed frames. We used the option of average-linkage that calculates the average distance between members of two clusters when minimum distance between clusters is greater than epsilon of 6.5.

### Inter-domain orientation calculation

To quantify the inter-domain motion between RRM1 and RRM2, we calculated the inter-domain orientation as described by Roca-Martínez et al. (14). Three conserved Cα positions were selected from each RRM domain: two on the β3 strand (RNP1) and one on the β1 strand (RNP2). From these, two vectors were defined per domain: the RNP1 vector, connecting the two β3 positions, and the RNP1–2 vector, connecting the third β3 position to the β1 position. Using the RRM1 vectors as normal vectors, two reference planes were defined: the RNP1 plane and the RNP1–2 plane. The corresponding RRM2 vectors were then normalized and projected onto these planes, and the angles between the projected RRM2 vectors and the RRM1 vectors were calculated using the arctangent function, resulting in two inter-domain angles with a full 360° range: the RNP1 angle (ψ), which captures the rotational component of the relative domain orientation, and the RNP1–2 angle (φ), which captures the displacement component. For the definition of vector points, we selected the following residues: from RRM1, A101 and A103 (RNP1) and I62 (RNP2); from RRM2, A181 and L183 (RNP1) and L139 (RNP2).

### Analysis of protein-RNA interface

Aromatic stacking interactions between protein aromatic residues and RNA bases were evaluated by monitoring two parameters. First, COM-COM distance was calculated between the ring atoms of each aromatic residue and the ring atoms of each RNA base. Second, the stacking angle (θ) was determined by computing the normal vector to each aromatic plane using the corrplane command in CPPTRAJ. The angle between the two normal vectors was calculated from the dot product of the vectors using the vectormath command. Together, these parameters allowed us to track the stability and geometry of the stacking interactions throughout the trajectory.

Hydrogen bonds analysis was performed in CPPTRAJ with a 3.5 Å distance cutoff and a 120° angle threshold, producing donor-to-acceptor hydrogen bond output files. These were then processed using an in-house script that aggregated the atom pair interactions into per-residue hydrogen bonds, assigning a count starting from 0 and incrementing for each atom pair interaction that matched the same residue pair. We considered separately two categories of hydrogen bonds formed by protein residues either with the RNA backbone or with the RNA bases (sequence specific). This produced per-residue hydrogen bond time series for the entire simulations, allowing us to track the formation and breaking of specific protein–RNA interactions and to distinguish between backbone and base-specific contacts.

We quantified the RNA backbone kink by defining two vectors from the phosphorus (P) atom of U6 to the phosphorus atoms of A4 and G8, respectively. The kink angle (Ω) was then calculated from the dot product between these two vectors using the vectormath function in CPPTRAJ. Using P atoms allowed us to measure backbone bending without the influence of ribose or base fluctuations.

## Results

### The DND1-RNA complex displays extensive conformational plasticity

The experimental structure of the DND1-RNA complex, determined by NMR spectroscopy (Fig 1A) shows 20 snapshots of the protein-RNA bound complex with structural differences confined to the N-terminal and C-terminal extensions (Fig. 1B).

**FIGURE 1.**
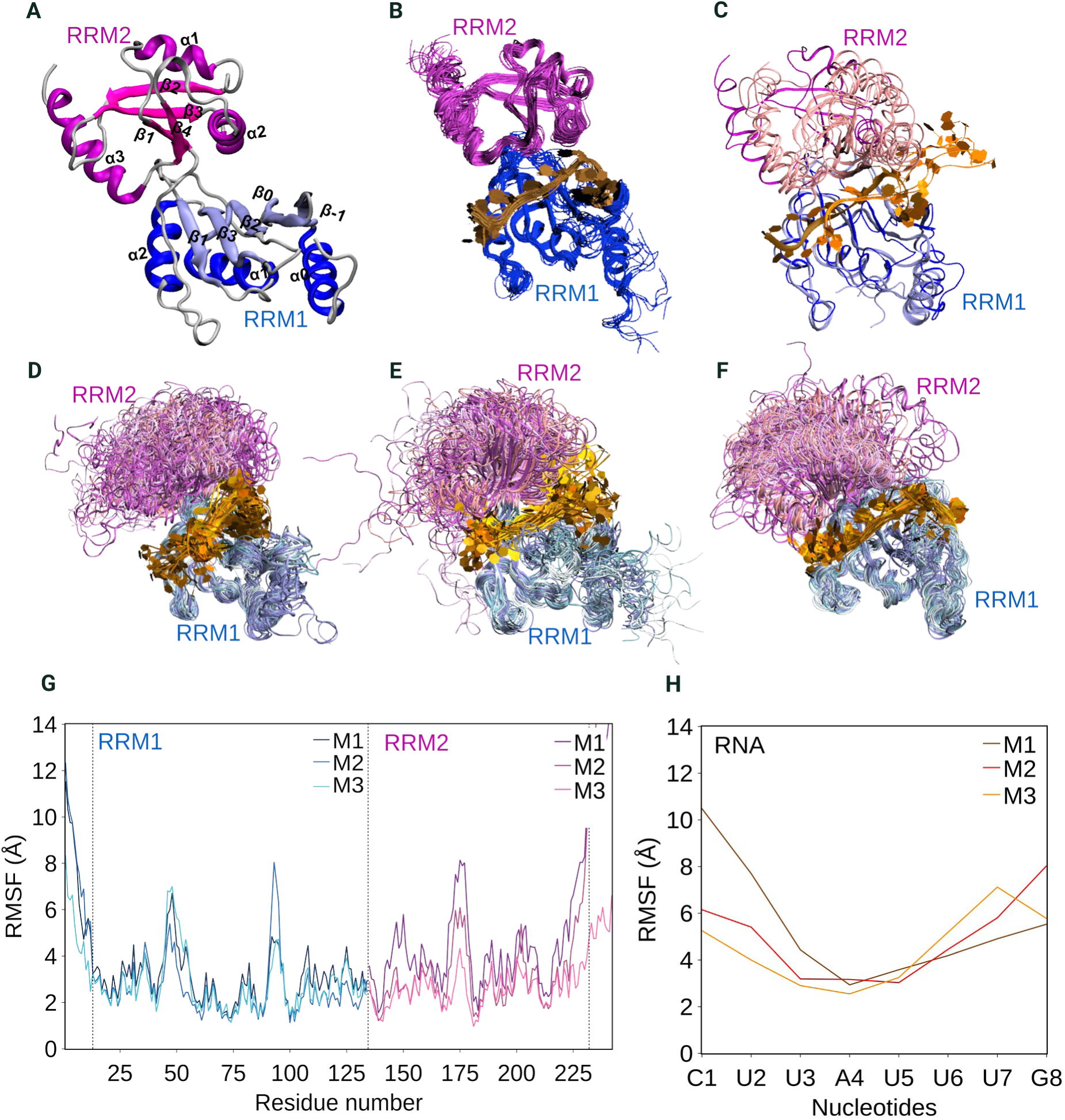
Flexibility of the DND1-RNA complex. (A) Average structure of the DND1 protein from the NMR ensemble (PDB-ID: 7Q4L). (B) The 20 structures of the experimental NMR ensemble superimposed. (C) 5 best AF3 models superimposed on the experimental structure (the frame nearest to the average structure of the NMR ensemble). Each model was the 1st ranked in each of the 5 runs with different random number seeds (see Methods). (D-F) Superimposed snapshots of the MD simulations taken every 500 ps from simulations M1 (D), M2 (E), and M3 (F). (G) RMSF of the protein atoms in the ensemble of the 3 MD simulations (RRM1 in blue, RRM2 in pink). (H). RMSF of the RNA atoms in the MD simulation ensemble. The colors in this figure will be kept throughout this manuscript unless indicated otherwise.

A comparison with 25 models generated by AlphaFold3 (AF3) resulted from 5 independent runs with different random number seeds, shows that these predicted structures are significantly different (Fig. 1C). In the AF3 models, the values for the fraction of native contacts (FNAT) at the binding site was from 0.018 to 0.38. These values combined with high global and interface RMSD (IRMSD) values confirmed that the predicted models differ significantly from the experimental structure (Table S3). However, the AF3 predicted structures represent only different predictions of the same structure and do not reflect conformational plasticity.

To investigate the dynamics of the complex, we performed a series of MD simulations (Table S2) and found that the complex samples many different conformations, most of them different from those found in the initial experimental structure (Fig. 1D-F, Fig. S1C). We observed large differences between the dynamics in the 3 replica simulations started with the model M2 (Fig. S1D). Therefore, we selected the replica (M2-R2) most similar to those started from the M1 and M3 models for further analysis. When mutating the central A4 nucleotide to U, we observed larger deviations from the reference structure (Fig. S1E), with the largest RMSD observed in the simulation started with the M2 model.

From residue-based RMSFs we found that the RRM1 domain is less flexible than the RRM2 domain in 2 out of 3 DND1-RNA simulations. In one simulation (M3), the RRM2 domain fluctuated less than in the other two (Fig. 1G). Investigating the flexibility of the RNA in the simulations, we found that the central adenine nucleotide in the RNA (A4), which is known from experimental data to be important for binding, was less flexible than other RNA nucleotides in all 3 DND1-RNA simulations (Fig. 1H). Therefore, our findings revealed that the DND1-RNA complex displays extensive conformational plasticity in the simulations, which explains the inability of AlphaFold3 to predict the conformation observed in the NMR structure.

When visualizing the simulations, we observed that flexibility is largely driven by changes in the orientation of the RRM2 domain relative to RRM1 which could explain the larger RMSF values for RRM2 (Fig. 1G). At the same time, the essential central adenine nucleotide in the RNA and to some extent the two flanking U bases remain more rigid (Fig. 1H). This finding is in agreement with the previously reported sequence-specific binding of DND1 to a UAU triad (18). Notably, the flexibility pattern was different in the two extra replica simulations started with the model M2 (Fig. S1F). When A4 was mutated to a U, the flexibility of the UAU central nucleotides in the RNA was increased, albeit U4 remained less flexible compared to its flanking nucleotides (Fig. S1G).

### RNA binding restricts the inter-domain motion of the two RRMs

To understand the nature of the conformational plasticity we observed, we first examined how the inter-domain orientation of the two RRMs changes during the simulations of the DND1-RNA complex and the free DND1. In the complex, the distance between the RRMs increased compared to the reference experimental structure in two of the three simulations, while in one simulation it remained in the interval observed in the NMR structure (Fig. 2A). To investigate whether the increase in the inter-domain distance was accompanied by changes in the inter-domain orientation, we first performed a hierarchical clustering on RMSD values calculated by fitting exclusively on RRM1 (Fig. S2A), which identified 17 distinct clusters of DND1-RNA conformations sampled during the simulations. Then, we evaluated the inter-domain orientation of each cluster representative structure by calculating the angles φ and ψ as described in the Methods. We found significant variation in the relative orientation of RRM2 relative to RRM1 compared to the NMR structure (Fig. 2B). We observed the largest variation along the φ angle. Superimposing the cluster representative structures on RRM1 revealed clear differences in how RRM2 was oriented relative to RRM1. For example, the structure of cluster C2 was similar to the reference except of a small rotation along ψ, while the structure in cluster C9 located at edge of the region of the φ-ψ conformational space which also contains the most populated clusters C0 and C1 showed a large displacement towards RRM1 (along φ) with little rotation, as seen from the orientation of the β1 strand in both structures (Fig. 2C).

**FIGURE 2.**
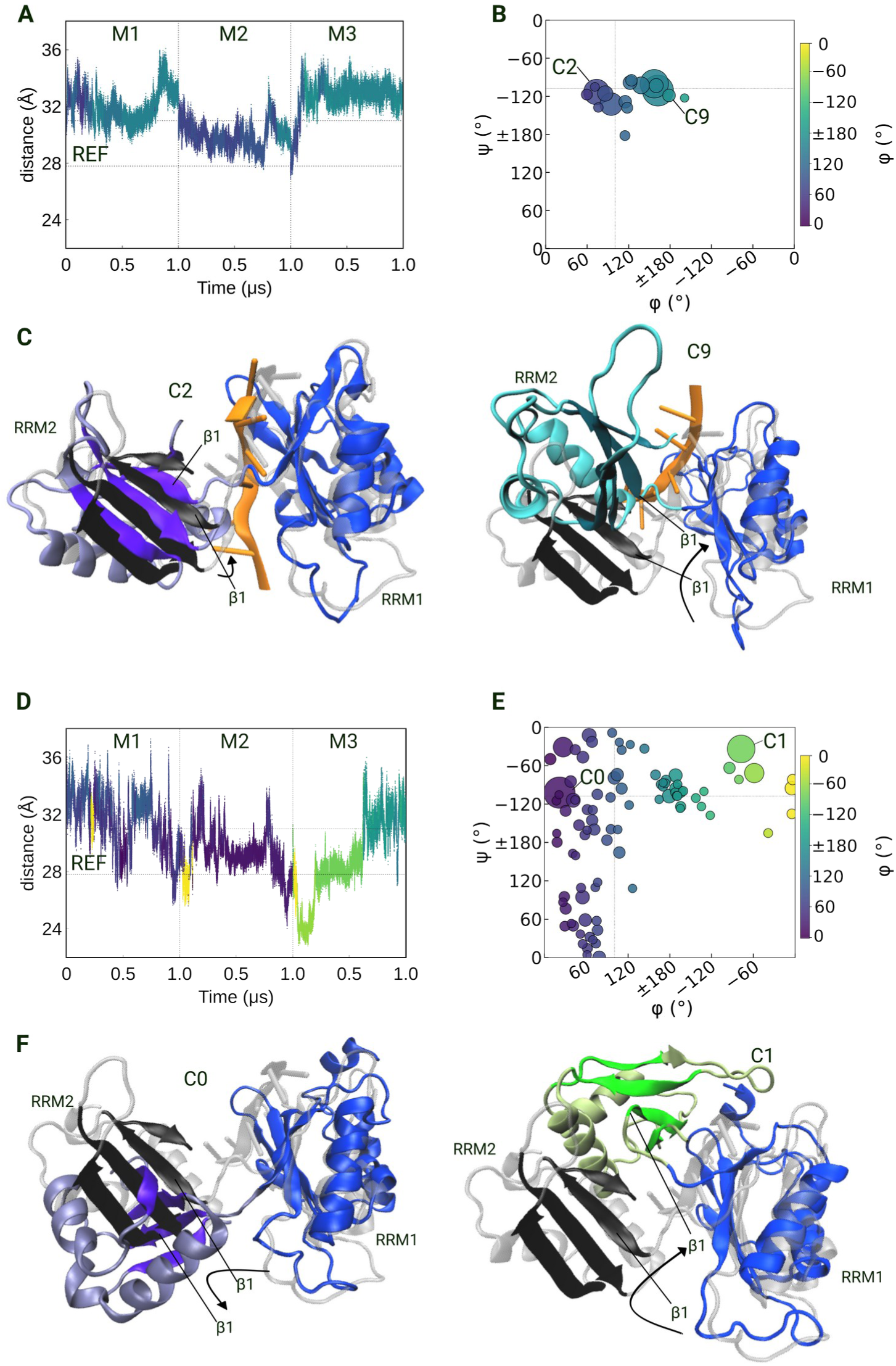
Inter-domain motion between RRM1 and RRM2. (A-C) DND1-RNA simulations. (D-F) DND1 simulations. Each panel is split into 2 rows. On the first row the RRM1-RRM2 center-of-mass distance during the 3 independent simulations (started with models M1, M2, and M3) is shown on the left and clusters of relative RRM1-RRM2 inter-domain orientations defined by angles φ and ψ are shown on the right. The angles were calculated according to Roca-Martinez et al (14) (see details in Methods). The clusters are colored by φ values, also mapped on the COM-COM distance plot. On the second row, selected cluster representatives superimposed on the reference experimental structure are shown. RRM1 is in blue, while the β-sheet of RRM2 is colored by the color of the cluster it represents (purple and green respectively). The rest of RRM2 is shown in slightly different nuances for a better view of the motion relative to the β-sheet. The reference structure is in gray with the β-sheet in black. For better visualization of the β-sheet orientations, we do not show the α3 helix of RRM2 in the structural snapshots.

In the absence of the RNA, the distance between the two RRMs varied significantly more (Fig. 2D). This increased inter-domain motion was also reflected in the hierarchical clustering (Fig. S2B), which produced 90 clusters, over five times more than in the RNA-bound system. The inter-domain orientation also spread over a wider range of the φ-ψ conformational space (Fig. 2E). The two most populated clusters illustrated the diversity of conformations sampled. While the representative structure of cluster C0 showed a displacement of RRM2 away from RRM1 (φ) accompanied by a small rotation, the representative structure of cluster C1 shows the opposite motion, with RRM2 displaced closer towards RRM1 (Fig. 2F).

Interestingly, when the central A4 was mutated in the RNA mutation increased RRM inter-domain flexibility and impacted the relative orientation of the 2 RRMs. This was clearly observed in the increased RMSDs of DND1 when superimposing only RRM1 (Fig. S2C). While the inter-domain distance remained near the interval observed in the NMR structure (Fig. S2D), there was a significant increase in the number of clusters obtained and in the spread of the inter-domain orientation angles (Fig. S2E).

Taken together, these results demonstrate that the DND1 system is highly dynamic, with the two domains exploring a broad conformational space with variable inter-domain orientations. However, the amplitude of these motions depends on the central A4 nucleotide.

### Cooperative inter RRM dynamics impact interface accessibility and restrict RNA motion

To understand how the inter-domain dynamics affect the accessibility of the main RNA binding interface and the RNA motion, we first measured the SASA of the RRM β-sheets (Fig. S3A-C) which is known to be the main RNA binding interface in the RRMs. We found an overall decrease in the SASA of the β-sheet of RRM1 in all systems during the simulations compared to the values in the experimental reference structure (Fig. 3A). The distributions were broader in the DND1-RNA and DND1 systems compared to the systems lacking RRM2 (Fig. 3A, S3D) reflecting the variability in the inter-domain orientation observed in these systems. Moreover, while there is a marginal increase of the RRM1 β-sheet SASA in the RRM1-RNA complex comparing the free RRM1, no significant difference was observed when comparing the DND1-RNA complex with the free DND1. The RRM1 β-sheet was the most accessible in the RRM1-RNA complex (Fig. 3A, S3D). In the case of RRM2, the accessibility of its non-canonical β-sheet varied significantly in the simulations of the DND1–RNA system. This complex sampled two distinct states, one with a higher SASA values for the RRM2 β-sheet near the upper limit of the values in the NMR structure and one with lower SASA values sampled mainly in the simulation started with the M3 model (Fig. 3B, S3C, E). Notable, with the exception of the DND1-RNA complex, the accessibility of the RRM2 β-sheet was similar in all simulations. These findings show that the absence of the RNA does not induce a collapsing of the two RRMs which could make the β-sheets inaccessible for RNA binding. The presence of the RRM2 and not the RNA binding has a compaction effect leading to a decrease in the RRM1 β-sheet SASA. At the same time, the accessibility of the RRM2 β-sheet is generally lower than that of RRM1 and is most sensitive to the inter-domain motions in the RNA-bound tandem RRM.

**FIGURE 3.**
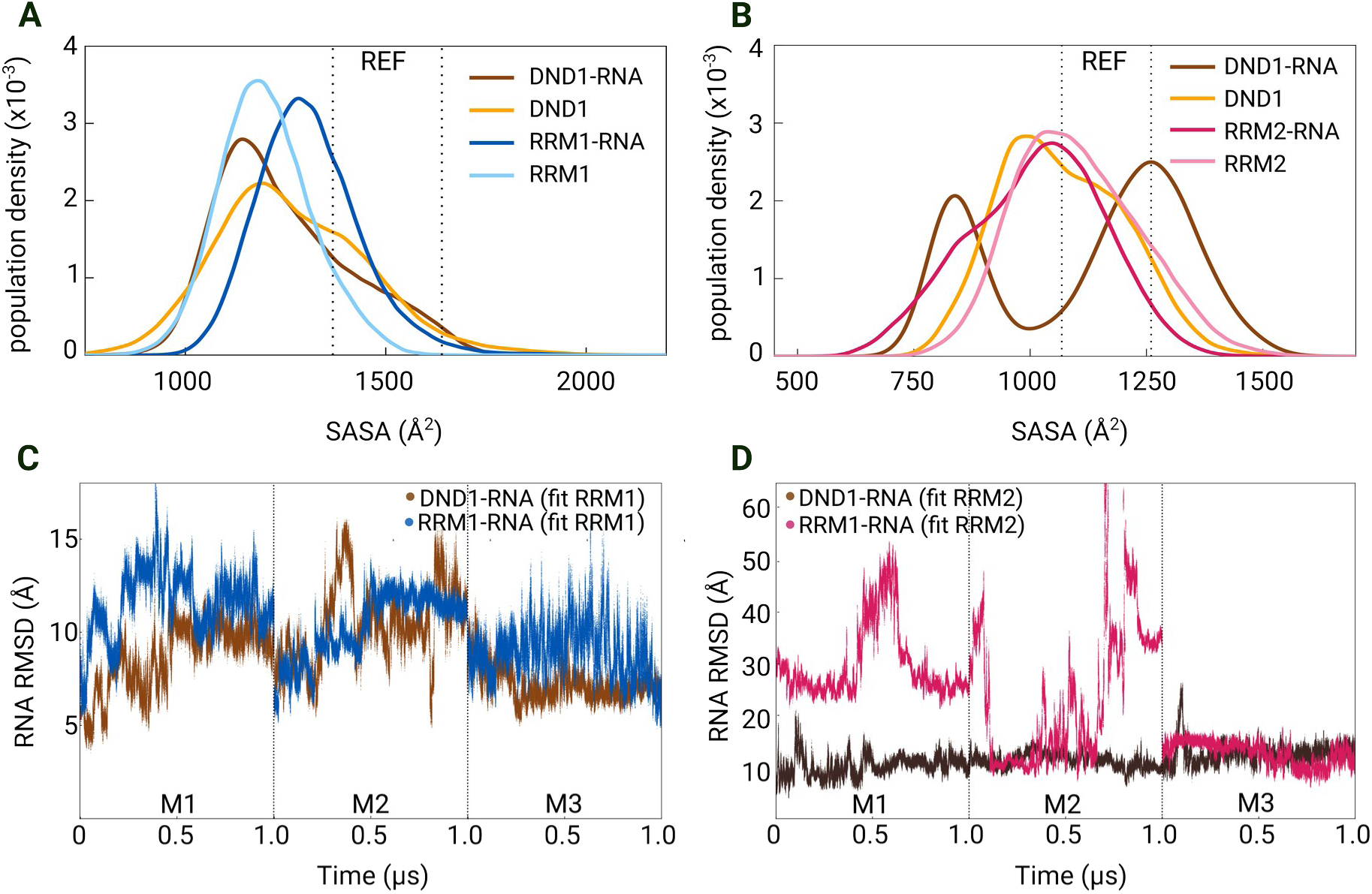
Tandem RRM cooperativity in RNA binding. (A-B) Accessibility of the β-sheet of RRM1 (A) and RRM2 (B) measured by the solvent accessible area (SASA). (C-D) RMSD of RNA when either RRM1 (C) or RRM2 (D) was fit to the reference structure. The DND1-RNA complex and DND1 protein are in brown and orange respectively. The individual domains are colored as in Figure 1. The dashed horizontal lines show the range of values in the reference experimental structure.

Next, we monitored how RNA motion is modulated by RRM binding either in tandem or in the absence of the second RRM by calculating the RMSD of the RNA while superposing either RRM domain. When we superimposed the trajectories on RRM1, the RNA deviated more from the reference structure when bound to RRM1 alone than when bound to both domains in tandem (Fig. 3C). Despite the difference observed, the RNA motion was significantly restricted by RRM1 binding even in the absence of RRM2. Only in one of the replica simulations started with the M2 model, we observed RNA unbinding in the RRM1-RNA complex (Fig. S3F). When aligning the trajectories on RRM2, the RNA deviated substantially more when bound to RRM2 alone (Fig. 3D, S3G). The RRM2-RNA binding did not restrict the motion of RNA in a similar manner observed for the RRM1-RNA binding. Only in the simulation of the RRM2-RNA complex started from the model M3, the RNA motion was restricted, with the RNA moving towards a more stable binding interface with the β-sheet of RRM2 (Fig. 3D). Interestingly, when we mutated the central A4 nucleotide, the motion of the RNA was significantly increased in the DND1-RNA^A4U^ complex but not in the RRM1-RNA^A4U^ (Fig. S3H).

### Tandem RRM binding favors a more open RNA conformation

To determine how protein binding affects the conformational space sampled by the RNA, we first calculated the ROG of the RNA. In the simulations of the free RNA (Fig. 4A), we observed continuous transitions between more compact and more extended structures, with a slightly higher population of more compact conformations with ROG values lower than those found in the experimental NMR structure (Fig. 4A). In contrast, in all protein-RNA complexes, the RNA sampled more extended conformations (Fig. 4A, Fig. S4A). In the DND1–RNA complex, the population of conformations with ROG values higher than in the reference structures was the largest. However, the RNA sampled some of the most compact conformations in the simulation of the DND1-RNA complex started with the M1 model and transitions between compact and extended conformations were sampled. Interestingly, in the truncated complexes with single RRMs, most sampled conformations had ROG values in the range of those found in the experimental structure of the tandem RRM-RNA complex. Nevertheless, we also observed several transitions from less compact to more compact conformations in these simulations (Fig. 4A). Structural snapshots corresponding to the peaks of the ROG distributions highlight these differences in the compactness of the sampled RNA conformations (Fig. 4B).

**FIGURE 4.**
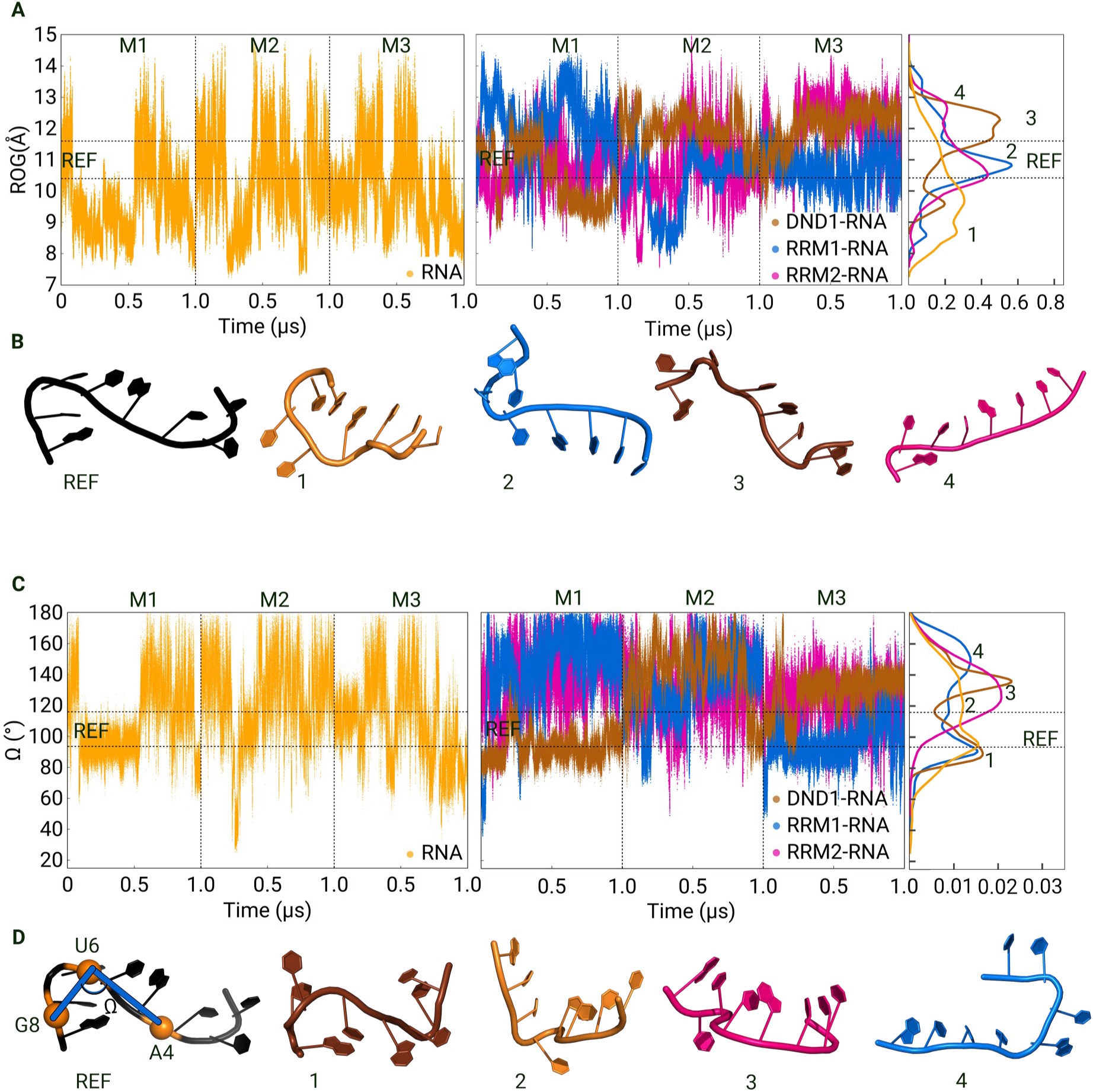
Conformational dynamics of the single-stranded RNA. (A) Radius of gyration (ROG) of the RNA during the 3 independent simulations (M1, M2, M3). The free RNA is shown in the left plot, while the bound RNA in the middle plot. On the right, histograms integrating the data from the 2 plots are shown (B) Representative structural snapshots selected from the peaks of the histograms (numbered) in (A). (C) The A4-U6-G8 kink angle in the RNA shown in the same sequence of plots as the ROG. (D) Representative structural snapshots selected from the peaks of the histograms (numbered) in (C). The colors are as in Figure 3 with the free RNA in yellow. The dashed horizontal lines show the range of values in the reference experimental structure.

In addition to the compactness, we also monitored the characteristic kink of the RNA conformation found in the NMR structure during the simulations (Fig. 4C,D). We observed that the kink angle (Ω) correlates well with the ROG in the free RNA, indicating that it is a good determinant of the RNA structure compactness. The free RNA sampled regular transitions between highly kinked and more extended conformations. Interestingly, in the simulations of the DND1-RNA complex Ω followed a distribution with two narrow peaks, one corresponding to a highly kinked structure similar to the most kinked conformation in the NMR ensemble sampled mainly in the simulation started with the M1 model and one less kinked conformation sampled mainly in the simulations started with the M2 and M3 models. The fluctuations of Ω were lower in the DND1-RNA complex, and we also observed a few transitions between the two types of conformations in the latter two simulations (Fig. 4C, Fig. S4B). In contrast, the truncated RRM-RNA complexes sampled wider distributions of Ω (Fig. 4C). Whereas in the RRM1-RNA complex, the RNA sampled the highly kinked conformation, in the RRM2-RNA complex the RNA was mostly found in highly variable more extended conformations. The alternative conformations sampled are illustrated with snapshots taken at the different peaks of the distribution histograms (Fig. 4D).

Interestingly, mutating the central A4 nucleotide into U had an important impact on the distribution of RNA conformations in the DND1-RNA complex (Fig. S4C, D), indicating that the conformation of the RNA bound to the tandem RRM is sensitive to the specific RNA sequence despite the overall plasticity observed in the simulations.

### Tandem RRM binding facilitates stable π-π interactions

To evaluate how the overall conformational stability described above modulates the sequence specific protein-RNA interactions, we analyzed the π-π stacking interactions and the hydrogen bonds between protein residues and RNA bases and backbone. Since aromatic π-π interactions play an important role in the stability of canonical RRM-RNA binding (3, 18), we examined whether these interactions were maintained in the DND1-RNA complex by monitoring the COM-COM distance between the interacting aromatic rings and the angle θ defined between the planes of the interacting rings (Fig. 5A). The central interaction is the π-π stacking of A4 on F61 in RRM1, and this remained stable throughout two out of three simulations (M1 and M3). In M2, A4 initially stacked on F61 in both parallel and T-shaped orientations, but towards the end of the simulation the base shifted its position to form a π-π interaction with Y102 which was maintained for the rest of the trajectory (Fig. 5B). In addition to this primary interaction, we found an additional π-π interaction between U5 with H189 in RRM2 which was also stable in 2 of the 3 simulations (Fig. 5C, Fig. S5A). We observed some variability in the stability of the stacking interactions between the 3 independent replicas started with the model M2 (Fig. S5B, C). Representative snapshots from the simulations illustrate these stacking orientations (Fig. 5D). Remarkably, the stacking interactions were much less present in the simulations of the RRM1-RNA complex (Fig. S5A). Moreover, there were no stacking interactions formed in the simulations of the RRM2-RNA complex. Finally, in the simulations with the mutated RNA^A4U^, U4 formed a stable interaction with F61 only in 1 of the 3 simulations (Fig. S5D–E). These findings highlight the role of the cooperative binding of both RRM in stabilizing these interactions. In addition, the findings reveal the importance of the identity of base A4 for forming stable stacking interactions in the DND1-RNA complex. These results demonstrated that the tandem arrangement of both RRM domains was required to sustain stable π-π interactions at the protein-RNA interface. While RRM1 provided the primary platform through F61, the cooperative presence of RRM2 was necessary to maintain these interactions over the course of the simulations.

**FIGURE 5.**
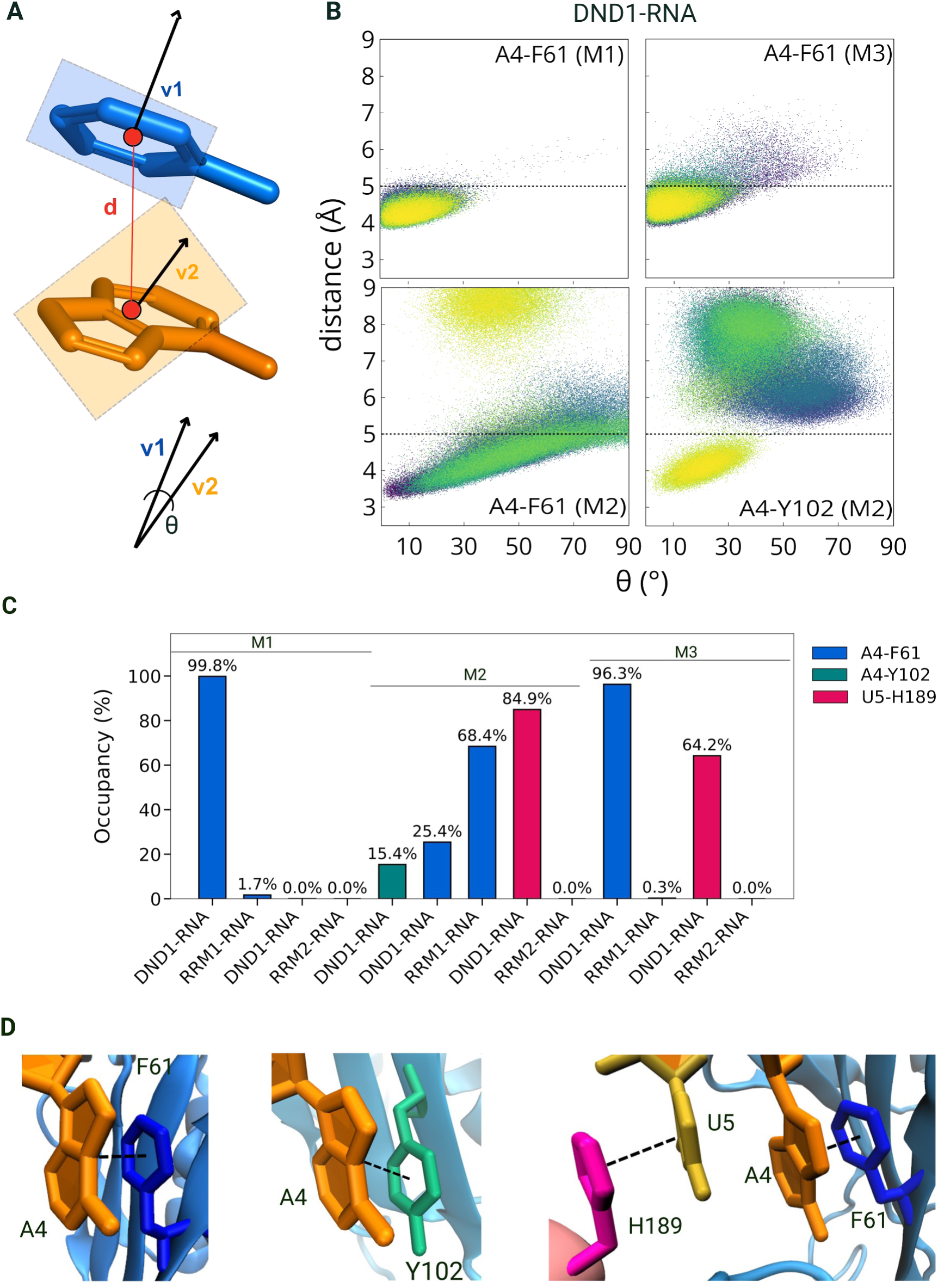
Stacking interactions of Adenine A4. (A) Schematic representation of the COM-COM distance and stacking angle (θ) definitions. (B) A4 stacking interactions in the DND1 system during the 3 independent simulations (M1, M2, and M3). Data points are colored by simulation time (µs). The horizontal dashed lines indicate the geometric thresholds defined for a stable stacking interaction. Parallel stacking corresponds to 0°<θ<45°, while T-shape stacking to 65°<θ<90°. (C) Occupancy of stacking interactions in each simulation. (D) Representative structural snapshots illustrating the interactions in (C).

### Core protein-RNA hydrogen bonds are maintained despite conformational plasticity

Overall, the number of protein-RNA contacts was higher and varied at least in the DND1-RNA complex with significantly more stable contacts established between RRM1 and RNA than between RRM2 and RNA (Fig. S6A). In the simulation of the truncated RRM2-RNA complex started from the M2 model, the RNA dissociated. Moreover, any observed increase of the number of RRM2-RNA contacts in the simulations of the RRM2-RNA complex was due to binding and re-binding of the RNA on different surfaces of RRM2. The same behavior was observed in the additional replica simulations started with the model M2 (Fig. S6B). Importantly, the number of protein-RNA contacts also decreased upon mutating A4 to a U, especially in the DND1-RNA complex (Fig. S6C).

In addition to the π-π stacking interactions analyzed above, the core network of hydrogen bonds between protein residues and nucleotides is also essential for the specificity of the DND1-RNA complex. Therefore, we also monitored the hydrogen bonds that were stable in more than 30% of at least one of the three simulations. We separated the hydrogen bonds in two categories: interactions with the RNA bases (sequence-specific) and with the RNA backbone (Fig. 6).

**FIGURE 6.**
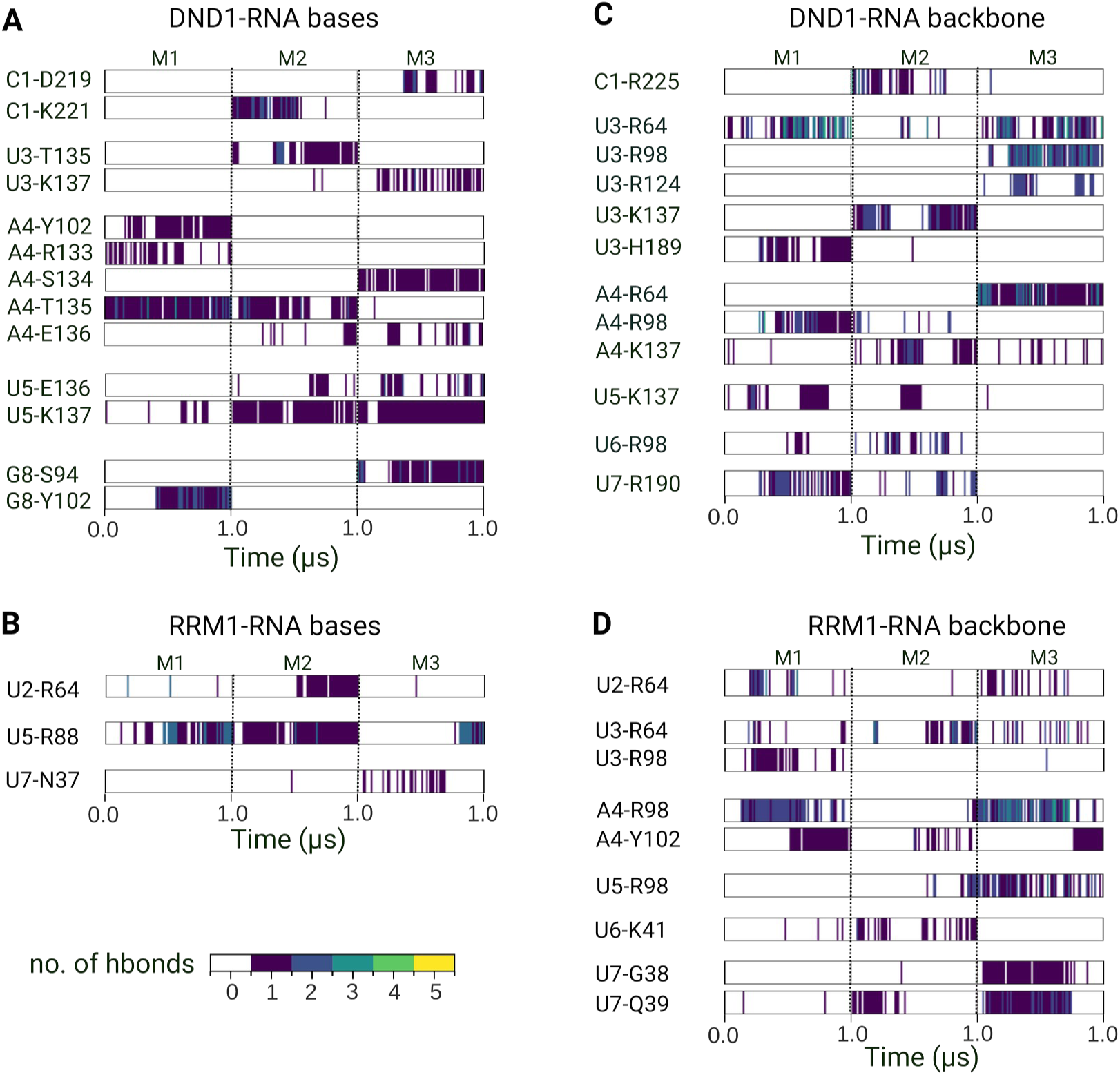
Hydrogen bonds at the protein–RNA binding interface. (A-B) Hydrogen bonds between protein and the RNA base in the DND1-RNA (A) and RRM1-RNA (B) systems. (C-D) Hydrogen bonds between the protein and RNA backbone in the DND1-RNA (C) and RRM1-RNA (D) systems. The occurrence of hydrogen bonds is shown during the 3 independent 1.0 µs simulations (M1, M2, M3). Only hydrogen bonds with an occupancy >30% in at least one of the three simulations are shown. The color bar (0–5) indicates the number of simultaneous hydrogen bonds detected per frame.

The sequence-specific interactions in the DND1-RNA complex were mainly formed by inter-domain linker residues. A4 formed stable hydrogen bonds either with S134 or T135. Some hydrogen bonds were also formed by A4 with R133 and E136, but these were generally restricted to small time intervals in a single simulation (Fig. 6A, Fig. S6D). The flanking nucleotides U3 and U5 also formed stable hydrogen bonds, in particular with K137 (Fig. 6A). In comparison, in the RRM1-RNA system only three hydrogen bonds involving RNA bases were present in more than 30% of at least one simulation. The most stable of these was the U5-R88 interaction (Fig. 6B).

In addition to sequence-specific hydrogen bonds, the RNA backbone also formed hydrogen bonds relevant for anchoring the RNA into the RRM binding site. In the DND1-RNA complex, R64 and R98 formed such hydrogen bonds with the backbone of U3 and A4 (Fig. 6C, Fig. S6E). Unlike the sequence-specific contacts, these arginine-mediated backbone interactions were also present in the RRM1-RNA complex, suggesting that the backbone anchoring was less dependent on the inter-domain linker and RRM2 orientation (Fig. 6D).

In the simulations with the mutated RNA^A4U^, both the base and the backbone of U4 formed significantly less hydrogen bonds with protein residues (Fig. S6F,G), providing additional evidence for the importance of A4 in the RNA recognition by DND1.

In conclusion, the protein-RNA interactions are mostly preserved in the DND1-RNA complex despite the overall conformational plasticity given by the inter-domain dynamics of the two RRMs. In the absence of RRM2, the key interactions are less conserved, whereas in the absence of RRM1, there is no stable protein-RNA interface in the MD simulations.

## Discussion

We report here that the DND1-RNA complex displays high conformational plasticity in MD simulations, deviating significantly from the experimentally determined structure (PDB-ID: 7Q4L) (18). The conformational variability is rooted mostly in the relative orientation of the two RRM motifs. The sampled conformations in the three independent simulations were different, suggesting that the simulation times are insufficient to achieve converged sampling of the inter-domain motion. Nevertheless, there were sufficient similarities between the three replicas per system which allowed us to explore and compare the dynamics observed in different systems. The simulations did show similarities in terms of protein-RNA interactions, especially for those that define the core specificity of DND1. This indicates that the variability observed among the independent simulations, particularly the changing in the relative orientation of the two RRM domains, showcases a dynamic protein-RNA interface which may allow multiple RRM-RRM arrangements while maintaining the core RNA binding features. Notably, AF3 also predicts a different tandem RRM-RNA binding configuration than that in the NMR structure, providing further evidence for the conformational plasticity of this complex.

This plasticity may be specific for tandem RRM motifs recognizing RNA in a non-canonical manner (18), without involving the main β-sheet surfaces of both RRMs in the interaction. Or it may depend on the structure and length of the linker connecting the two RRMs which differ in tandem RRM containing proteins (14). While the conformational flexibility of tandem RRMs is often linked to long, unstructured inter-domain linkers (e.g., in SRSF1 binding;(39)), DND1 presents a structural paradox. It features a notably short, four-residue inter-domain linker, and its C-terminal RRM2 is non-canonical yet still crucial for RNA binding. As a key regulator, DND1 must bind long RNA sequences and facilitate protein-protein interactions (19). Our findings show that the free DND1 protein inherently explores a wide range of conformations, driven by the flexibility of the tandem RRM arrangement. Upon RNA binding, this conformational freedom is substantially narrowed, though the two domains still retain a baseline level of mobility within a restricted orientational range. Notably, the A4U mutation partially disrupts this structural restraint, revealing that the identity of the central nucleotide is directly involved in stabilizing the relative inter-domain orientation.

Our simulations highlight that the DND1-RNA system maintains significant conformational plasticity despite the short linker and non-canonical RRM2. This inherent flexibility may be involved in its biological role, enabling DND1 to adapt its structure to various interactions. This finding is supported by a new crystal structure of the ternary DND1-RNA-NANOS3 complex (preprint 40) which presents a different inter-domain orientation of the two RRMs compared to the original DND1-RNA complex structure (PDB-ID: 7Q4L) (18). In light of this evidence, our findings underscore that the conformational plasticity of tandem RRMs is not solely dependent on canonical structure or linker length, but is a fundamental mechanism allowing DND1 to adapt to binding other proteins in diverse regulatory processes. The mechanism of high-affinity RNA recognition by DND1 relies on the cooperative binding of its tandem RRMs. Previous studies demonstrated that the maximum binding affinity is achieved only when both RRM domains are engaged (18, 41). In the case of DND1’s cooperative model, the RRM1 domain functions as the main binding interface, while the RRM2 domain serves as a stabilizing anchor. Our findings confirm the cooperative mechanism with RRM1 maintaining a partially stable interface with the RNA even in isolation, while RRM2 acting as a cooperative partner that cannot sustain a stable interaction on its own. When both domains bind RNA in tandem, the conformational space sampled by the RNA is more restricted than in the complexes with either RRM domain, demonstrating that the cooperative binding of both RRMs is required to constrain the RNA in the complex.

Based on our simulations, we propose a layered binding model for the DND1-RNA interaction. The first layer is the π-π stacking of RNA bases on protein aromatic residues, primarily A4 on F61, which provides the foundation for a stable interaction. The simulations where both A4 and U5 stacking were present (on F61 and H189, respectively) further demonstrated the importance of engaging both RRMs in tandem, and the ability of A4 to switch stacking partners from F61 to Y102 in the M2 simulation suggests that the aromatic surface of RRM1 can accommodate alternative binding geometries while maintaining the core interaction. This is supported by experimental data (18), showing that mutants F61A and Y102A could not be purified or expressed in soluble form, and suggesting that these aromatic residues are critical not only for RNA binding but also for the structural integrity of the protein. The second layer consists of the arginine-mediated backbone contacts, primarily through R64 and R98, which anchor the RNA on its binding surface. Consistently, the R98A mutation in RRM1 was shown to completely abolish RNA binding (18), highlighting the importance of these contacts. The third layer involves the hydrogen bonds with the inter-domain linker residues, which position and stabilize the RNA bases within the binding pocket. The combination of these three layers: aromatic stacking, backbone anchoring, and linker-mediated hydrogen bonds, together creates the stable and specific DND1-RNA interaction.

In our simulations of the truncated RRM1-RNA complex, the RNA binding remained partially stable even in the absence of RRM2, confirming RRM1’s intrinsic capacity for high-affinity recognition. However, compared to the tandem DND1-RNA system, the RRM1-RNA simulations showed fewer stable hydrogen bonds and a lower overall number of contacts, indicating that while RRM1 can sustain binding on its own, the full set of interactions is only achieved when RRM2 is present. This underscores RRM1’s role as the initial, flexible docking site that requires the cooperative contribution of RRM2 to form the complete binding interface.

The RRM2 domain, while contributing to the final stability, likely functions primarily as a cooperative trap for the RNA. Previous mutational studies targeting the “unusual RNA-binding pocket” of RRM2 (K197A and W215F variants) resulted in significantly weaker binding, but did not completely abolish it, suggesting that these single mutations only partially disrupt the pocket (18). In our DND1-RNA simulations, we observed in 2 out of 3 simulations a stable π-π interaction between H189 and U5, located on the α-helix 2 of RRM2. This is particularly interesting as it mirrors the non-canonical RNA recognition mode described for the SRSF1 pseudo-RRM, where conserved residues on the α-helix rather than the β-sheet mediate RNA binding (42). However, in most of the RRM2-RNA simulations, stable contacts and hydrogen bonds were not maintained, and two simulations showed episodes of complete RNA unbinding. This aligns with experimental data showing that RRM2 alone does not have detectable binding to RNA (18) and supports a model where RRM2 cannot independently sustain a stable complex but instead increases the overall binding affinity when acting in tandem with RRM1.

In such a dynamic mechanism, it is plausible that the interaction between DND1 and its target RNA proceeds via a hybrid mechanism, combining elements of conformational selection and induced fit. The initial engagement is likely achieved through conformational selection, as the RRM1 domain binds using a pre-established conformation to an RNA conformation which is also sampled by the free RNA. Then the RRM2 domain reorients and converges towards RRM1 via an induced fit mechanism, forming an enclosure mechanism that traps the RNA and locks the domains into a stable complex.

The accuracy with which the simulations can predict such a mechanism depends on the force field parameters used. Historically, RNA force fields suffered inaccuracies in describing structural ensembles of single-stranded RNA. Conventional non-polarizable force field models like AMBER and CHARMM often overestimate electrostatic forces. This can lead to overly stable and compact complexes in the simulation, which may mask genuine conformational plasticity. While polarizable models may offer increased accuracy, and AMOEBA is demonstrated as a viable option for studying protein-RNA systems (43), Baltrukevich and Bartos ultimately conclude that no single force field consistently outperforms others, suggesting no perfect solution for MD force field selection (44).

The accurate modeling of RNA is fundamentally challenged by its high conformational flexibility and requires precise force field parameterization to prevent computational artifacts such as unintended structural deviations and the formation of non-native loop conformations (45–48). This difficulty is underscored by the existence of advanced force field alternatives, such as those incorporating NBfix corrections specifically designed to mitigate the collapse of single-stranded RNA (49). We used the AMBER OL3 force field (34) for RNA coupled with the 4-point OPC (Optimal Point Charge) water model (32). This combination is widely regarded as a robust and reliable benchmark for the simulation of nucleic acids (48–50). Interestingly, with this combination we did not observe the collapsing of the RNA into a compact structure as reported previously (51, 52).

The ionic conditions were carefully chosen to reproduce the protocol used to generate the NMR structure (18) which did not contain Mg^2+^. While the inclusion of Mg^2+^ ions is frequently considered in RNA simulations (53, 54), accurately modeling the properties of divalent ions remains inherently difficult. For this reason, Mg^2+^ inclusion is generally recommended only when its binding site is definitively known (48). We added both Na+ and K+ monovalent ions to ensure that no ion is overconcentrated in the highly negatively charged systems and to reflect the physiological presence of both ion types and create a diversity in the sizes and properties of ions that surround the molecules.

Previous MD simulation studies on protein-RNA interactions consistently highlight the inherent challenge of accurately modeling these highly dynamic systems and the flexibility of their binding mechanisms. Krepl et al. combined over 50 μs of MD simulations with NMR data on the Fox-1 and SRSF1 RRM-RNA complexes and showed that the method is robust enough to reliably describe the structural dynamics of RRM-RNA interfaces, while also revealing that several segments of the interface involve competition between dynamical substates rather than firmly formed interactions (55). More recently, the same group used millisecond-scale unbiased simulations to capture spontaneous binding of single-stranded RNA to RRM domains, demonstrating that binding proceeds through pre-binding states and dynamic scanning of the RNA sequence (56), which is consistent with the hybrid conformational selection and induced fit mechanism we propose for DND1.

Increased flexibility and a broader range of binding modes were also observed in MD simulations of the single-domain FUS protein in complex with a stem-loop RNA structure (57). Similarly, an MD study examining the tandem RRM protein TDP-43 binding RNA showed a stable complex, but with differences in the binding mode compared to experimental data (58) These examples underscore the critical point that MD simulations are complementary to structural experimental techniques like NMR or X-ray crystallography. By revealing the dynamic mechanism of these interactions, the simulations bring essential new insights not available from static experimental structures.

Additional MD simulations with different protocols may help in the thorough assessment of the force fields for such dynamic RNA based systems. Moreover, MD simulations of a tandem RRM proteins with canonical and non-canonical binding patterns in the future could provide a robust comparison platform for decoding the dynamics of tandem RRM-RNA interactions. This could open the door for future design of RRM containing proteins with special RNA binding features and for targeting tandem RRM proteins in disease.

## Data Availability Statement

The MD simulations data with input and output files are provided at https://zenodo.org/records/17940554, https://zenodo.org/records/17940953, and https://zenodo.org/records/17944396

## Author contribution

RGV and VC designed and performed the molecular dynamics simulations, analysed the data, and wrote the manuscript

## Acknowledgments

The Cojocaru group (https://cojocarulab.eu) is part of the Doctoral School in Integrative Biology of the Babes-Bolyai University. RGV is funded by the ministry of education of Romania. We thank Alexandre Bonvin and the Computational Structural Biology group at Utrecht University (https://bonvinlab.org) for support, feed-back and discussion. Vlad Cojocaru thanks Hans Schöler for continuous support and the Max Planck Computing and Data Facility (https://www.mpcdf.mpg.de/) for computer resources.

## Declaration of generative AI and AI-assisted technologies in the manuscript preparation process

During the preparation of this work, the author(s) used Claude for obtaining an initial list of correctly formatted references. The author(s) reviewed and edited the output as needed and take full responsibility for the content of the published article.

