## Supplementary Information for "Conformational plasticity and sequence specificity interplay in non-canonical tandem RRM-RNA binding"

[2]Computational Structural Biochemistry Group, STAR-UBB Institute, Babeş-Bolyai University, Cluj-Napoca, Romania

Short title: Non-canonical tandem RRM-RNA dynamics

Keywords: RRM domain; RNA-binding proteins; conformational change; sequence specificity; molecular recognition; tandem RRM; molecular dynamics simulation

* Correspondence to:

Vlad Cojocaru

Computational Structural Biochemistry Group

STAR-UBB Institute, Babeş-Bolyai University

Tiberiu Popoviciu street 2-4, Cluj-Napoca, Romania

Web: https://cojocarulab.eu

**Table of Contents**

**Supplementary Figures……………………………………………………………..………..3**

**Figure S1.** Flexibility of DND1-RNA complexes…………....…………..……………..………..…3

**Figure S2.** Interdomain dynamics in the DND1-RNA complexes……….…………..………..…..….…4

**Figure S3.** RRMs interface accessibility and RNA dynamics relative to each RRM……………...…....5

**Figure S4.** RNA compactness and kink angle dynamics..…....………….………………….………..6

**Figure S5.** π–π stacking interactions across simulations……………………………...……..…..….7

**Figure S6.** Hydrogen bonding and contact analysis…………….....................................…..………..8

**Supplementary Tables………………………………………………………………….….9**

**Table S1.** Equilibration parameters …………………………………………….....…….….………...9

**Table S2.** Summary of simulation systems..........................................................................………......10

**Table S3.** HADDOCK3 caprieval scoring of AlphaFold3-predicted DND1–RNA models.…………...11

The MD simulations data: input files, scripts for the equilibration steps and production runs, log files and output files are provided at 10.5281/zenodo.17940953, 10.5281/zenodo.17940554, 10.5281/zenodo. 17944396.

### Supplementary Figures

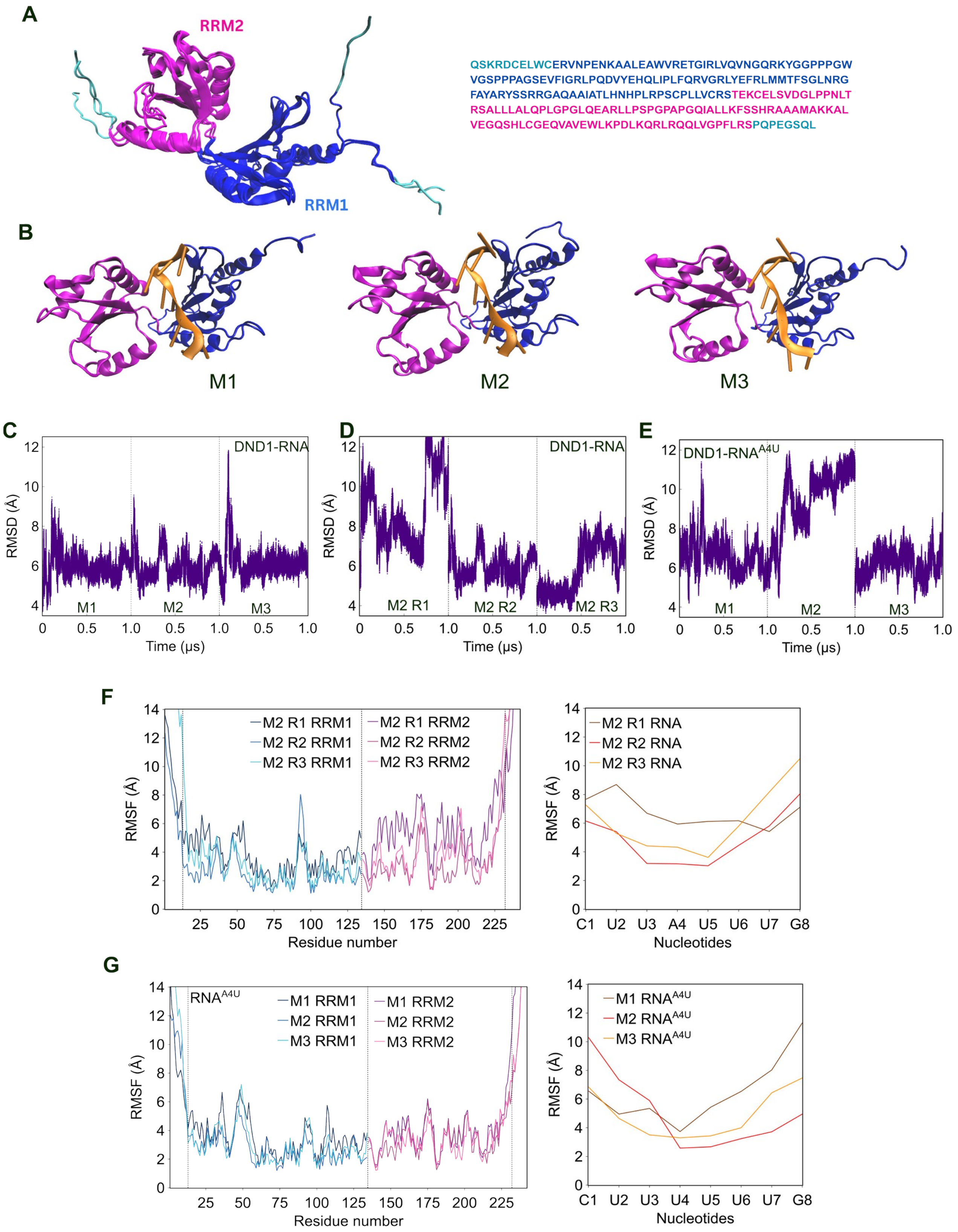

**Figure S1. Structure and flexibility of the DND1-RNA complex.** (A) The NMR structural ensemble (pdb id: 7Q4L) with added N-terminal and C-terminal tails (cyan) (left) and the DND1 sequence (right). (B) NMR structures (frames 1, 8, and 13 from the ensemble corresponding to models M1, M2, and M3) used as starting structures for MD simulations (see Methods for details). (C-E) RMSD calculated by fitting all protein heavy atoms to the reference structure of DND1-RNA in the simulations included in the main article (C), the M2-R1, M2-R2, M2-R3 (D), and the DND1-RNA$^{A4U}$ simulations (E). (F) RMSFs in the 3 independent replicas (R1, R2, and R3) started with the M2 model. (G) RMSFs in the 3 independent simulations of the DND1-RNA$^{A4U}$ complex started from models M1, M2, and M3. See also Figure 1.

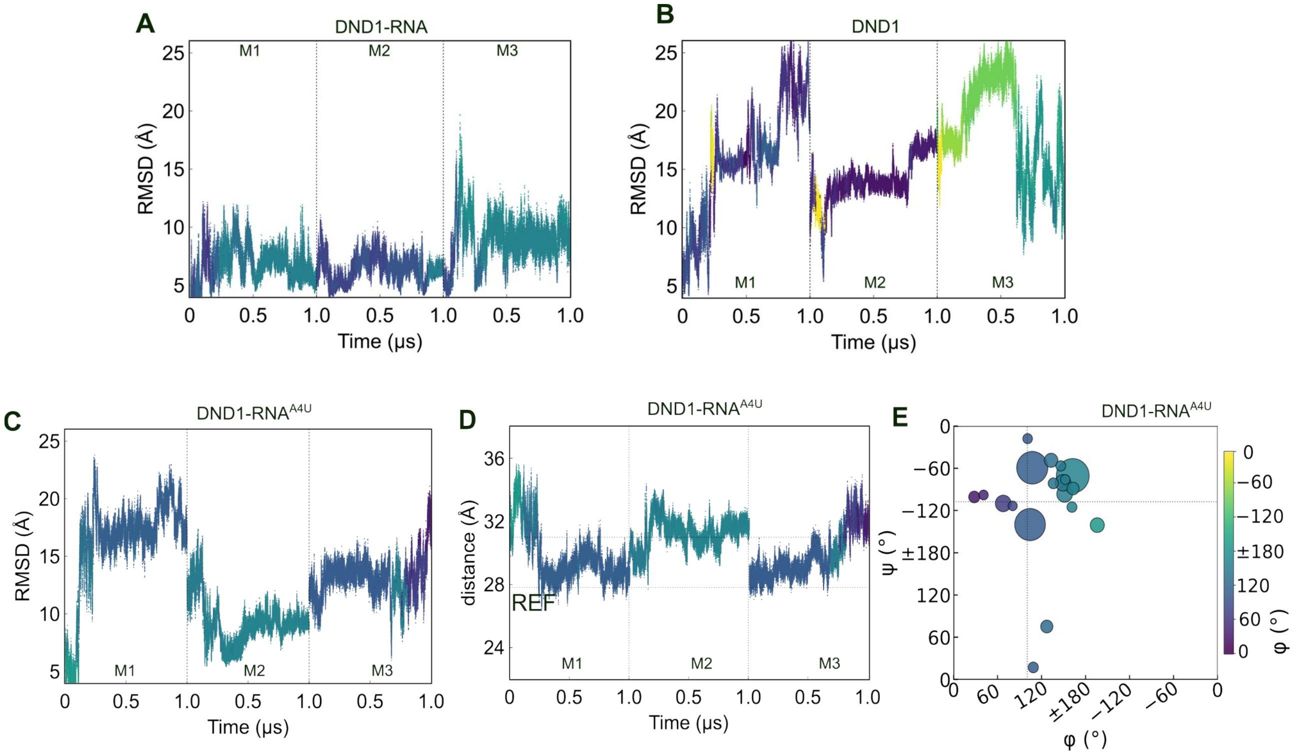


**Figure S2. Interdomain dynamics.** (A-B) RMSD of the DND1 after fitting only RRM1(used for the clustering analysis) for DND1-RNA (A) and (B) DND1 systems. (C) RMSD as in (A) during the DND1-RNA$^{A4U}$ simulations. (D) Time evolution of the inter center-of-mass (COM) distance between RRM1 and RRM2 in the DND1-RNA$^{A4U}$ simulations. Data is color-coded by the orientation angle $\varphi$. (E) Interdomain conformational landscape from the clustering analysis (see Methods for details). See also Figure 2.

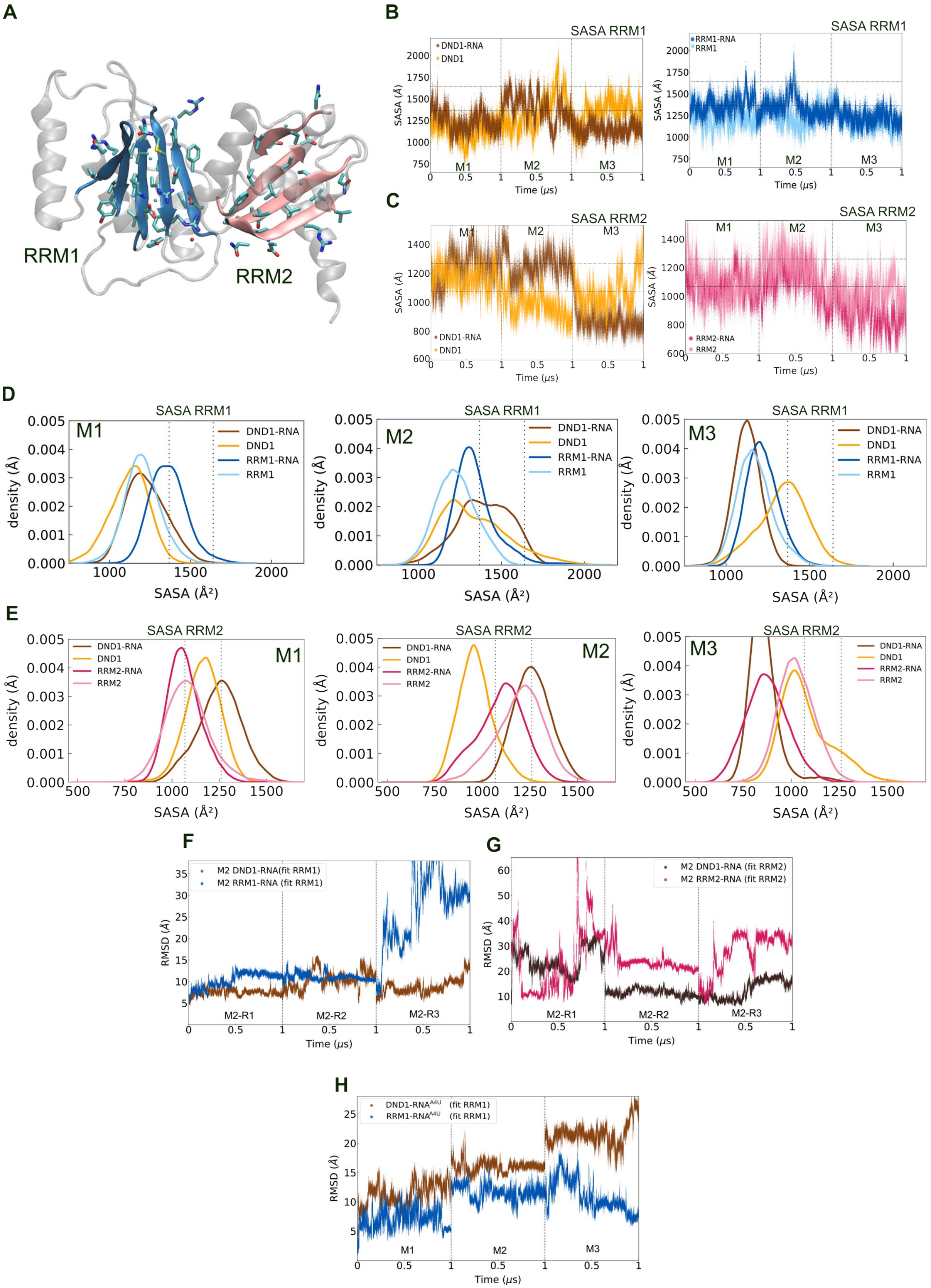

A
RRM1
RRM2
B
SASA RRM1
DND1-RNA
DND1
RRM1-RNA
RRM1
M1
M2
M3
Time (μs)
SASA (Å)
C
SASA RRM2
RRM2-RNA
RRM2
D
E
density (Å)
SASA (Å²)
F
M2 DND1-RNA(fit RRM1)
M2 RRM1-RNA (fit RRM1)
M2-R1
M2-R2
M2-R3
RMSD (Å)
G
M2 DND1-RNA (fit RRM2)
M2 RRM2-RNA (fit RRM2)
H
DND1-RNA$^{A4U}$ (fit RRM1)
RRM1-RNA$^{A4U}$ (fit RRM1)

**Figure S3. Accessibility and dynamics at the RRM1-RRM2 interface.** (A) Structural overview of the β-sheet residues selected for SASA calculations in RRM1 (blue) and RRM2 (pink). (B-C) SASA time series for RRM1 (B) and RRM2 (C). (D-E) SASA density distributions for RRM1 (D) and RRM2 (E). (F –G) RMSD of the RNA calculated after fitting only the RRM1 (F) or RRM2 (G) in the simulations with the original RNA. (H) RMSD of the RNA calculated after fitting only the RRM1 in the simulations with the A4U mutation in the RNA. See also Figure 3.

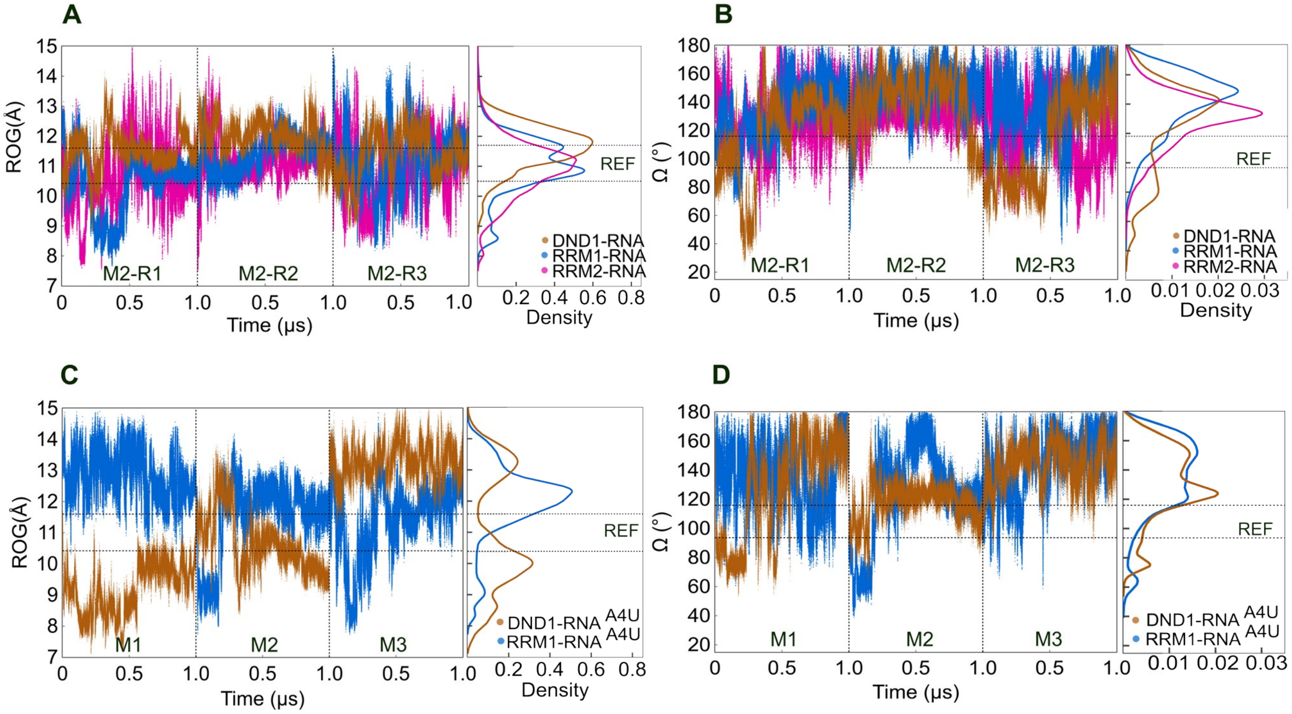


**Figure S4. RNA compactness and kink angle.** (A-B) Radius of gyration (ROG) of the RNA (A) and the RNA kink angle (Ω) (B) in the simulations with the original RNA. (C-D) Radius of gyration (ROG) of the RNA$^{A4U}$ (C) and the RNA kink angle (Ω) (D) in the simulations with the A4U mutated RNA. DND1–RNA complex is in brown), RRM1-RNA in blue, RRM2-RNA in magenta. Each panel shows the time series on the left and the corresponding histograms on the right. Dashed lines indicate the reference values from the NMR structure. See also Figure 4.

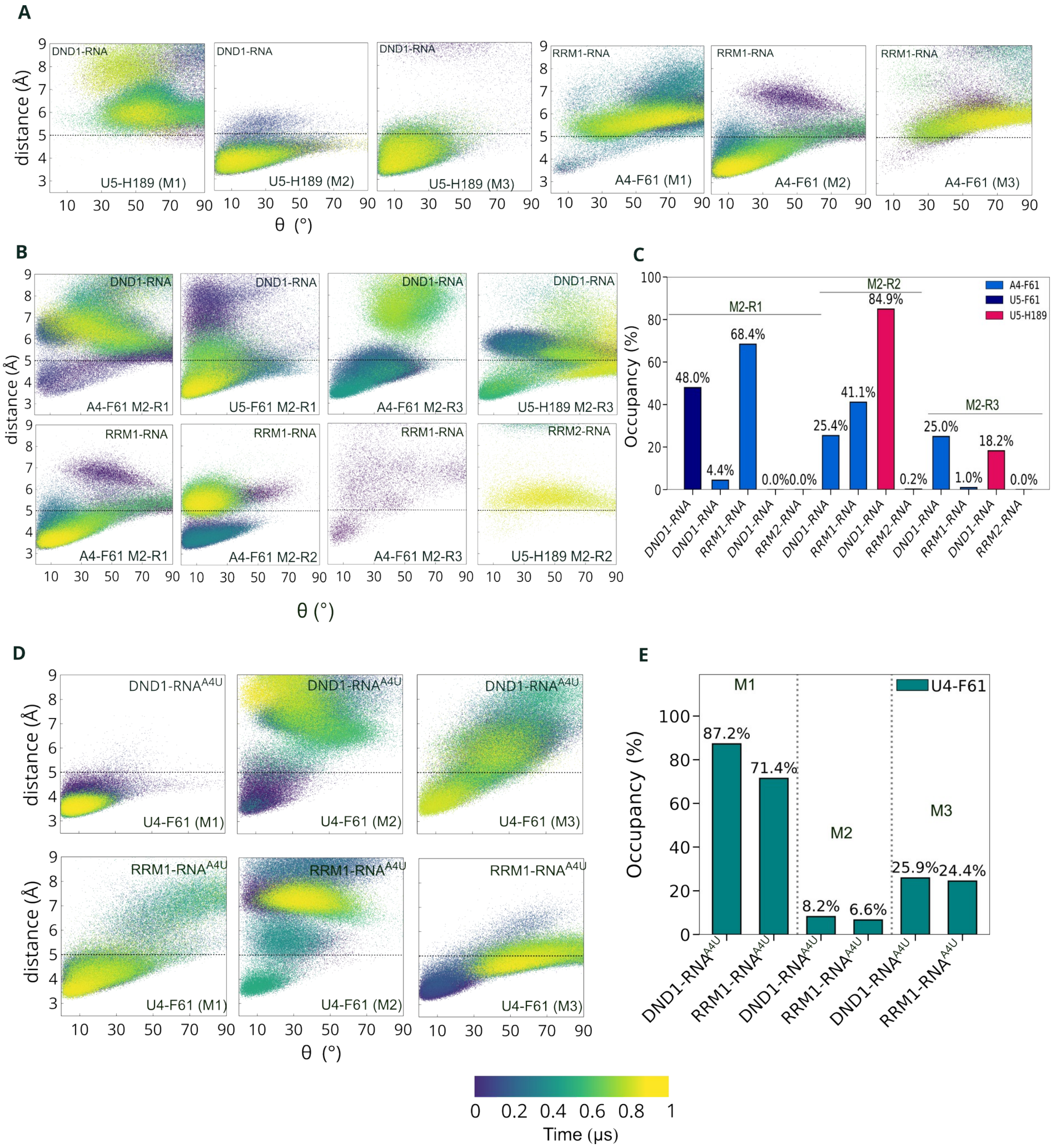


**Figure S5. π–π stacking interactions.** (A) Distance and θ angle between conjugated pairs of aromatic rings from the protein and the RNA in the simulations included in the main manuscript (M1, M2, M3) colored by simulation time. (B) Same as in (A) but for the 3 independent replica simulations started with model M2. (C) Occupancy of stacking interactions shown in (B). (C) Same as in (A) and (B) but for the simulations with the A4U mutated RNA. (E) Occupancy of stacking interactions shown in (D). See also Figure 5.

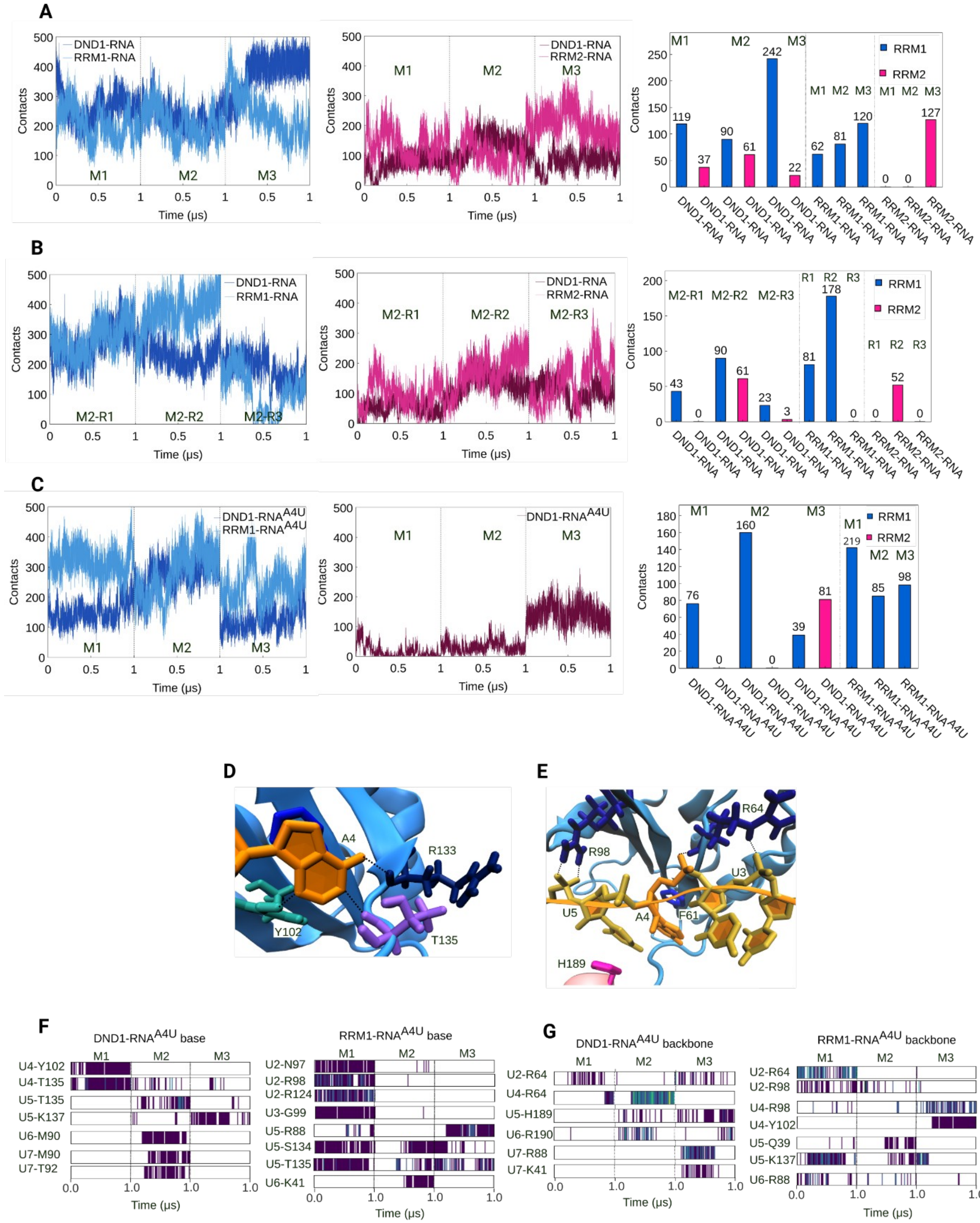


**Figure S6. Contacts and hydrogen bonds analysis.** (A) Total number or contacts between RRM1 (left, dark blue for DND1-RNA complex and light blue for the RRM1-RNA complex) or RRM2 (middle, magenta for DND1-RNA complex and pink for the RRM2-RNA complex) and the number of stable RRM-RNA contacts (occupancy >50%, same coloring). (B) Same as in (A) but for the 3 independent replica simulations started with model M2. (C) The same as in (A) and (B) but for the simulations with the A4U mutated RNA. (D) Representative hydrogen bonds between A4 and protein residues R133, T135, and Y102 in the simulation started with the M1 model. (E) Arginine-mediated hydrogen bonds (R64, R98) with the U3–A4–U5 backbone in the M3 replica. Dashed lines indicate hydrogen bonds. (F) Time series of RNA base–protein hydrogen bonds with occupancy exceeding 30% in at least one simulation for the DND1–$RNA^{A4U}$ and RRM1–$RNA^{A4U}$ systems. (G) As in (F), for RNA backbone–protein hydrogen bonds.

**Supplementary Tables**

**Table S1. Equilibration parameters**. Multi-stage equilibration protocol used for the simulations detailing the gradual reduction of positional restraints and adjustment of MD parameters (e.g., timestep, damping, constraint scaling) across 17 sequential steps.

| Step | timestep | constraints | Constraint Scaling | Langevin Damping | ensemble | Restraints type | extraBonds |
|---|---|---|---|---|---|---|---|
| eq01 | 1 | on | 25 | 5 | NVT | all solute restrained | off |
| eq02 | 1 | on | 10 | 1 | NPT | structured part protein-RNA complex | off |
| eq03 | 1 | on | 10 | 1 | NPT | structured part protein-RNA complex | off |
| eq04 | 1 | on | 5 | 1 | NPT | structured part protein-RNA complex | off |
| eq05 | 1 | on | 5 | 1 | NPT | structured protein-RNA + restraints to keep the interface stable | on |
| eq06 | 1 | on | 1 | 1 | NPT | structured protein-RNA + restraints to keep the interface stable | on |
| eq07 | 1 | on | 1 | 1 | NPT | structured protein-RNA + restraints to keep the interface stable | on |
| eq08 | 1 | on | 0.1 | 1 | NPT | structured protein-RNA + restraints to keep the interface stable | on |
| eq09 | 1 | on | 0.1 | 1 | NPT | structured protein-RNA + restraints to keep the interface stable | on |
| eq010 | 1 | off | | 1 | NPT | restraints to keep the interface stable | on |
| eq011 | 1 | off | | 1 | NPT | restraints to keep the interface stable | on |
| eq012 | 1 | off | | 1 | NPT | restraints to keep the interface stable | on |
| eq013 | 1 | off | | 1 | NPT | restraints to keep the interface stable | on |
| eq014 | 1 | off | | 1 | NPT | | off |
| eq015 | 1.5 | off | | 1 | NPT | | off |
| eq016 | 2 | off | | 1 | NPT | | off |
| eq017 | 2 | off | | 0.1 | NPT | | off |

**Table S2. Summary of simulation systems.** Each system was initiated from three NMR-derived conformers (M1, M2, M3) of the experimental structure (PDB: 7Q4L). Additional M2 replicas were performed after sequence-specific interactions broke early in the first replica. $RNA^{A4U}$ mutant simulations were included to probe the stability of A4 sequence-specific contacts.

| M1 | simulation time | M2 | simulation time | M3 | simulation time |
|---|---|---|---|---|---|
| DND1-RNA | 1 μs | DND1-RNA | 1 μs | DND1-RNA | 1 μs |
| DND1 | 1 μs | DND1 | 1 μs | DND1 | 1 μs |
| RRM1-RNA | 1 μs | RRM1-RNA | 1 μs | RRM1-RNA | 1 μs |
| RRM2-RNA | 1 μs | RRM2-RNA | 1 μs | RRM2-RNA | 1 μs |
| RRM1 | 1 μs | RRM1 | 1 μs | RRM1 | 1 μs |
| RRM2 | 1 μs | RRM2 | 1 μs | RRM2 | 1 μs |
| RNA | 1 μs | RNA | 1 μs | RNA | 1 μs |

| M2 R2 | simulation time | M2 R3 | simulation time |
|---|---|---|---|
| DND1-RNA | 1 μs | DND1-RNA | 1 μs |
| RRM1-RNA | 1 μs | RRM1-RNA | 1 μs |
| RRM2-RNA | 1 μs | RRM2-RNA | 1 μs |

| M1 | simulation time | M2 | simulation time | M3 | simulation time |
|---|---|---|---|---|---|
| $DND1\text{-}RNA^{A4U}$ | 1 μs | $DND1\text{-}RNA^{A4U}$ | 1 μs | $DND1\text{-}RNA^{A4U}$ | 1 μs |
| $RRM1\text{-}RNA^{A4U}$ | 1 μs | $RRM1\text{-}RNA^{A4U}$ | 1 μs | $RRM1\text{-}RNA^{A4U}$ | 1 μs |

**Table S3. HADDOCK3 caprieval scoring of AlphaFold3-predicted DND1–RNA models.** All 25 models generated across 5 AF3 webserver runs (test tx, seed number) ranked by HADDOCK3 caprieval scores. RMSD: all heavy atoms of the full RRM–RNA complex relative to the experimental NMR reference. IRMSD: heavy atoms at the RRM–RNA interface only. Gray-highlighted rows indicate the top-ranked models by AF3 pLDDT, shown in Figure 1C.

| AF3 model | Caprieval rank | rmsd | irmsd | fnat | lrmsd | ilrmsd | dockq |
|---|---|---|---|---|---|---|---|
| fold_dnd1_t1_50_model_0 | 1 | 6.424 | 5.989 | 0.18 | 12.982 | 12.391 | 0.18 |
| fold_dnd1_t1_50_model_1 | 2 | 6.884 | 6.02 | 0.22 | 14.026 | 12.314 | 0.182 |
| fold_dnd1_t3_150_model_0 | 3 | 6.594 | 6.5 | 0.08 | 14.961 | 13.87 | 0.125 |
| fold_dnd1_t3_150_model_1 | 4 | 6.33 | 6.316 | 0.1 | 14.139 | 13.4 | 0.14 |
| fold_dnd1_t3_150_model_2 | 5 | 6.866 | 6.184 | 0.18 | 14.355 | 13.155 | 0.165 |
| fold_dnd1_t3_150_model_3 | 6 | 6.244 | 5.294 | 0.38 | 11.278 | 10.243 | 0.272 |
| fold_dnd1_t3_150_model_4 | 7 | 5.815 | 5.071 | 0.36 | 11.207 | 10.166 | 0.269 |
| fold_dnd1_t4_200_model_0 | 8 | 6.153 | 5.814 | 0.08 | 12.946 | 12.171 | 0.148 |
| fold_dnd1_t4_200_model_1 | 9 | 6.393 | 5.852 | 0.34 | 12.739 | 11.587 | 0.237 |
| fold_dnd1_t4_200_model_2 | 10 | 6.961 | 6.276 | 0.14 | 14.195 | 12.996 | 0.153 |
| fold_dnd1_t4_200_model_3 | 11 | 6.303 | 5.354 | 0.36 | 11.456 | 10.623 | 0.263 |
| fold_dnd1_t4_200_model_4 | 12 | 6.115 | 5.458 | 0.32 | 12.439 | 11.179 | 0.236 |
| fold_dnd1_t1_50_model_2 | 13 | 7.14 | 6.39 | 0.14 | 14.634 | 13.386 | 0.148 |
| fold_dnd1_t5_250_model_0 | 14 | 6.633 | 6.339 | 0.08 | 14.432 | 13.317 | 0.13 |
| fold_dnd1_t5_250_model_1 | 15 | 6.752 | 6.545 | 0.08 | 15.891 | 14.423 | 0.117 |
| fold_dnd1_t5_250_model_2 | 16 | 5.756 | 5.784 | 0.14 | 13 | 12.306 | 0.167 |

| | | | | | | | |
|---|---|---|---|---|---|---|---|
| fold_dnd1_t5_250_model_3 | 17 | 5.076 | 4.596 | 0.42 | 9.653 | 8.967 | 0.318 |
| fold_dnd1_t5_250_model_4 | 18 | 6.709 | 6.019 | 0.14 | 13.94 | 12.799 | 0.156 |
| fold_dnd1_t1_50_model_3 | 19 | 6.197 | 5.986 | 0.42 | 13.49 | 12.469 | 0.254 |
| fold_dnd1_t1_50_model_4 | 20 | 6.318 | 4.994 | 0.46 | 10.768 | 9.48 | 0.309 |
| fold_dnd1_t2_100_model_0 | 21 | 7.069 | 6.597 | 0.16 | 15.572 | 14.149 | 0.146 |
| fold_dnd1_t2_100_model_1 | 22 | 6.38 | 5.916 | 0.2 | 13.258 | 12.314 | 0.184 |
| fold_dnd1_t2_100_model_2 | 23 | 6.289 | 5.224 | 0.44 | 11.54 | 10.038 | 0.289 |
| fold_dnd1_t2_100_model_3 | 24 | 6.231 | 5.062 | 0.36 | 12.074 | 10.48 | 0.257 |
| fold_dnd1_t2_100_model_4 | 25 | 6.388 | 4.556 | 0.3 | 9.69 | 8.398 | 0.278 |
